# The role of MEGF10 in myoblast fusion and hypertrophic response to overload of skeletal muscle

**DOI:** 10.1101/2024.10.19.619219

**Authors:** Louise Richardson, Ruth Hughes, Colin A Johnson, Stuart Egginton, Michelle Peckham

## Abstract

Biallelic mutations in multiple EGF domain protein 10 (*MEGF10*) gene cause EMARDD (early myopathy, areflexia, respiratory distress and dysphagia) in humans, a severe recessive myopathy, associated with reduced numbers of PAX7 positive satellite cells. To better understand the role of MEGF10 in satellite cells, we overexpressed human MEGF10 in mouse *H-2k*^*b*^-tsA58 myoblasts and found that it inhibited fusion. Addition of purified extracellular domains of human MEGF10, with (ECD) or without (EGF) the N-terminal EMI domain to *H-2k*^*b*^-tsA58 myoblasts, showed that the ECD was more effective at reducing myoblast adhesion and fusion by day 7 of differentiation, yet promoted adhesion of myoblasts to non-adhesive surfaces, highlighting the importance of the EMI domain in these behaviours. We additionally tested the role of Megf10 *in vivo* using transgenic mice with reduced (*Megf10*^+/-^) or no (*Megf10*^-/-^) Megf10. We found that the extensor digitorum longus muscle had fewer Pax7 positive satellite cell nuclei and was less able to undergo hypertrophy in response to muscle overload concomitant with a lower level of satellite cell activation. Taken together, our data suggest that MEGF10 may promote satellite cell adhesion and survival and prevent premature fusion helping to explain its role in EMARDD.

## Introduction

Myopathies, diseases of skeletal muscle, impair the ability of a muscle to regenerate in response to damage, either directly affecting muscle fibre activity or indirectly through effects on muscle stem (satellite) cells. In the recessive congenital myopathy, EMARDD (early myopathy, areflexia, respiratory distress and dysphagia), the ability of muscle to regenerate is impaired, but the reason for this is still unclear. Skeletal muscle fibres in EMARDD patients have a reduced diameter, fewer nuclei per fibre and lack Pax7^+^ satellite cells (Logan et al., 2011). The disorder is caused by mutations in *MEGF10* (multiple epidermal growth factor-like domains 10). MEGF10, the membrane protein encoded by this gene, has been suggested to be important for satellite cell interaction with the extracellular matrix (Logan et al., 2011). Mutations in *MEGF10* that cause EMARDD appear to reduce proliferation and migration of activated satellite cells, resulting in fewer myogenic cells that can eventually fuse together to form new adult myofibres (Holterman et al., 2007; Li et al., 2021; Saha et al., 2017). MEGF10 has also been suggested to promote satellite cell proliferation, whilst regulating myogenic differentiation (Holterman et al., 2007).

In addition to its role in satellite cells, MEGF10 has been reported to be required for engulfment, a similar role to that reported for the *C. elegans* CED-1 protein (Callebaut et al., 2003; Holterman et al., 2007; Suzuki and Nakayama, 2007b), an orthologue of MEGF10. Exogenous expression of MEGF10 in Hela cells induces these cells to engulf the protein GULP (Hamon et al., 2006). This role in engulfment is particularly important in the brain, where MEGF10 is highly expressed in astrocytes and likely plays a role in synapse trimming (Park and Chung, 2023). More recently MEGF10 has been found to be enriched in neuromuscular junctions with a possible role in modifying these synapses (Juros et al., 2024).

MEGF10 has a large extracellular domain that comprises an N-terminal EMI domain followed by 17 epidermal growth factor (EGF)-like domains (Fig. 1A). The EMI domain was first described in proteins in the EMILIN family of glycoproteins and is associated with protein multimer formation (Colombatti et al., 2011). The EMI domain of MEGF10, whilst having the conserved consensus sequence at the C-terminus (WRCCPG(Y/F)xGxxC), has only 6 rather than the 7 cysteine residues found in many EMI domains, and thus is more similar to the EMI domain in multimerin. This domain has been predicted to interact with the membrane phospholipid phosphatidylserine (PS), which is exposed on the surface of apoptotic cells as a signal marking the cell for phagocytosis *via* TTR-52 (Tung et al., 2013), consistent with the potential role of MEGF10 in engulfment. Interestingly, PS is also exposed on the surface of skeletal muscle myoblasts during fusion (Jeong and Conboy, 2011; van den Eijnde et al., 2001) and the PS receptor BAL1 promotes myoblast fusion (Hochreiter-Hufford et al., 2013). Thus, the EMI domain of MEGF10 could also play a role in myoblast-myoblast adhesion in fusing cells, via PS.

**Figure 1:**
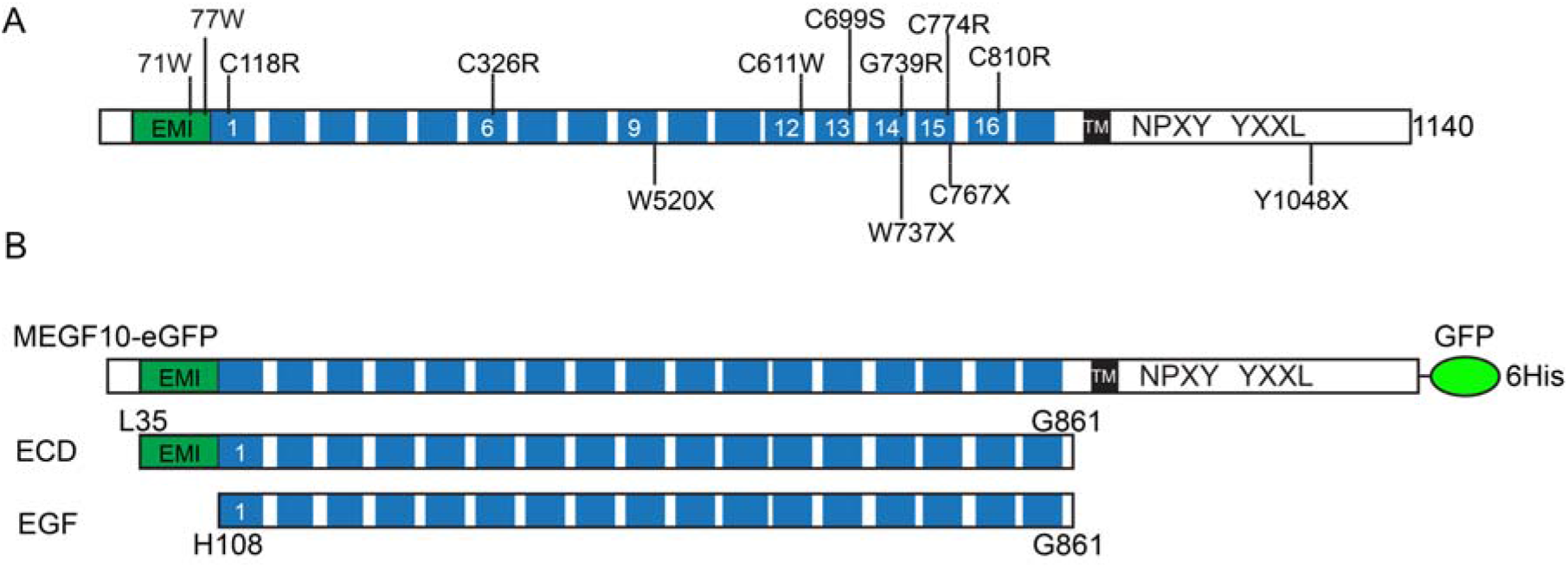
MEGF10 domains, and constructs used in the *in vitro* experiments. A: The domains of MEGF10, showing positions of known disease mutations (Fujii et al., 2022). B. The three constructs used in the *in vitro* experiments: MEGF10-eGFP, in which eGFP is fused to the C-terminus of MEGF10; and the ECD and EGF domains, expressed and purified using a mammalian cell system (see methods).

The EGF domains of MEGF10 comprise two different forms. There are 12 EGF-like domains composed of 7 conserved cysteine rich residues and 5 laminin-type EGF-domains that contain 8 conserved cysteine residues capable of forming four disulphide bonds. EGF domains may have a role in mediating intercellular signalling and act in receptor-ligand interactions (Wouters et al., 2005). Thus, these extracellular domains mediate a role for MEGF10 in cell adhesion (Suzuki and Nakayama, 2007a).

Satellite cells can contribute to skeletal muscle hypertrophy, in which skeletal muscle fibre diameter increases in response to hormonal, endocrine or mechanical stimuli, resulting in increased girth and strength of the muscle (reviewed in (Bagley et al., 2023). Hypertrophy commonly results from activities such as resistance training or mechanical overload. It results in transient increases of rapamycin complex 1 (mTORC1), which increases muscle protein synthesis, but increases in satellite cell number and in myonuclear accretion can also be important (Roberts et al., 2023). Mechanical overload models in mice and rats can mimic this response. One type of overload model involves surgery to remove the *tibialis anterior* (TA) muscle from one leg, which forces the synergistic *extensor digitorum longus* (EDL) to undergo sustained stretch, and thereby work harder when the animal is ambulant (Egginton et al., 2011). It results in a mild hypertrophy compared to unloaded muscle. This type of muscle perturbation is more physiological than inducing acute muscle damage by injecting toxins (Mahdy, 2019). The muscle overload model is also useful in determining how disease states may affect muscle hypertrophy. Previous work using the *mdx* mouse, which is a model for Duchenne muscular dystrophy (DMD), has shown that overloaded EDL muscle is unable to withstand the added strain resulting from the removal of the TA muscle, and undergoes accelerated deterioration (Dick and Vrbova, 1993).

The exact roles and function of MEGF10 in skeletal muscle are still poorly understood. Here, we have explored the role of MEGF10 in myoblast adhesion and fusion *in vitro* using a myoblast clone from the *H-2k*^*b*^-tsA58 transgenic mouse (Morgan et al., 1994; Richardson et al., 2022). We tested the effects of overexpressing MEGF10 on fusion and migration. We further tested the effects of MEGF10 by adding expressed and purified exogenous MEGF10 domains to cultured myoblasts to determine effects on cell adhesion, migration and fusion. We then explored the role of Megf10 in hypertrophy and satellite cell activation in response to overload *in vivo*, using a knockout mouse for *Megf10*. Use of the muscle overload model indicates that in humans, MEGF10 has a potential role in satellite cell function that would blunt the overload response, indicating a potential role in the aetiology of EMARDD.

## Results

### Overexpression of EGFP-MEGF10 reduces fusion and cell motility of cultured myoblasts

To explore MEGF10 effect on myoblast differentiation *in vitro*, we used a single *H-2k*^*b*^-tsA58 myoblast clone (C1F) derived from satellite cells (Morgan et al., 1994; Richardson et al., 2022). We generated an adenovirus to express either eGFP or eGFP-fused to the C-terminus of MEGF10 (Fig. 1B). The resultant constructs were the expected size (Supplemental Fig. 2). Tests for the optimal MOI (multiplicity of infection) demonstrated that an MOI of ∼100 was optimal (Supplemental Fig. 2).

**Figure 2:**
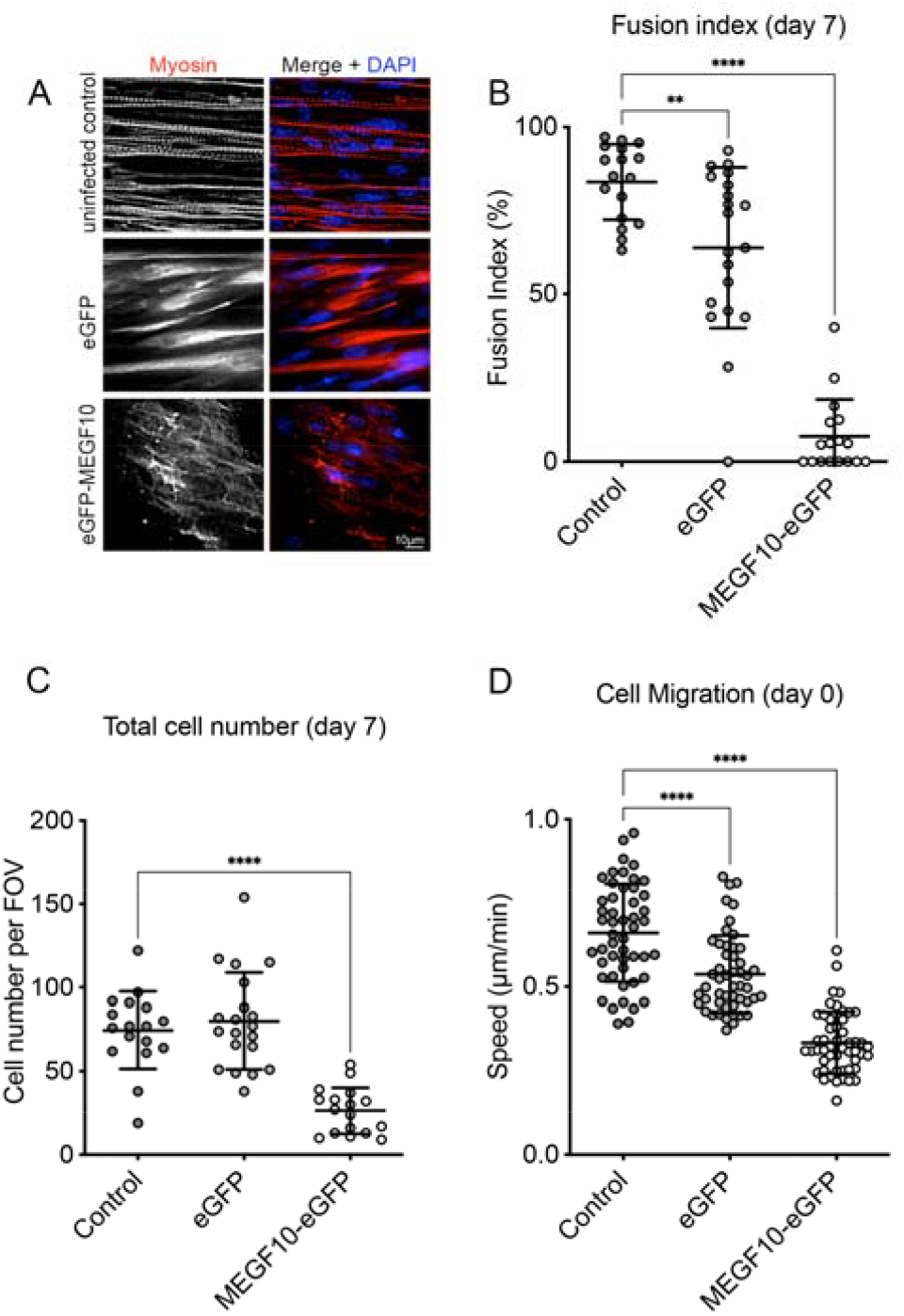
Effects of overexpression of MEGF10-eGFP on fusion, cell number and migration. **A**: Images of fused cells at day 7 for myoblasts (non-infected cells), myoblasts infected with an eGFP expressing adenovirus (MOI 100) and myoblasts infected with MEGF10-eGFP expressing adenovirus (MOI 100). Myotubes were stained for skeletal myosin (red) and DAPI (blue). **B:** Analysis of fusion using a minimum of 5 fields of view from 3 separate experiments. Fusion index was measured at day 7 (differentiation conditions). **C**: total number of cells present at day 7. **D:** Cell migration measured after 24 hours of expression. Data shows multiple individual measurements from three biological replicates. The mean ± the standard deviation (S.D.) is shown together with the results from an ANNOVA. ** *P*<0.01, **** *P*<0.001

Uninfected wild type C1F myoblasts fused into multinucleated myotubes with high efficiency (fusion index of ∼80%, Fig. 2A, B) when cultured under differentiation conditions. Expression of eGFP, using the adenovirus, significantly reduced fusion to 64% (Fig. 2A, B), suggesting that viral infection alone may reduce fusion. However, expression of MEGF10-eGFP reduced the fusion index to very low levels (fusion index of 7.5%, Fig. 2A, B). Thus, overexpression of MEGF10 inhibits fusion.

A reduction in myoblast fusion could arise from effects on cell viability or proliferation resulting from MEGF10□eGFP expression. We found that expression of MEGF10-eGFP significantly reduced cell number at day 7 of differentiation (Fig. 2C). This effect is unlikely to be due solely to infection with the adenovirus, as expression of GFP alone using an adenoviral construct did not affect cell number. Thus, expression of MEGF10-eGFP is likely to reduce cell proliferation or could also affect cell viability. Expression of MEGF10-eGFP also significantly reduced myoblast motility compared to cells expressing eGFP and uninfected cells (Fig. 2D). This reduction in migration may also contribute to the reduction in fusion.

### The extracellular domain of MEGF10 reduces fusion of cultured myoblasts

Imaging of C1F myoblasts infected with adenovirus to express MEGF10-eGFP at different MOIs (Supplemental Fig. 2B) showed that the higher the MOI, the more MEGF10-eGFP was localised to the Golgi. At the MOI of 100 used in these experiments, MEGF10-eGFP was localised to intracellular vesicles and Golgi and plasma membrane, and thus we cannot rule out that intracellular as well as membrane localised MEGF10-EGFP contributes to the effects we observe on cell fusion. To rule out intracellular effects, we carried out additional experiments in which we tested the effects of adding expressed and purified MEGF10 extracellular domain constructs to cultured myoblasts. We generated two constructs, an extracellular domain comprising all of the EGF domains together with the N-terminal EMI domain (ECD) and a shorter extracellular domain that lacks the N-terminal EMI domain and thus comprises the EGF domains only (EGF).

Pure MEGF10 extracellular domain (ECD) and EGF domain (lacking EMI domain) (Fig. 1B), was obtained by expression and purification of these constructs in HEK293 cells (Supplemental Fig. 3A, B). The molecular weights of these proteins were higher than predicted (Supplemental Fig. 4A, B); the molecular weights of both ECD and EGF were found to be approximately 130kDa, whereas the predicted molecular weights are 91kDa and 81kDa, respectively. We confirmed that the expressed and purified protein were the correct MEGF10 constructs by western blotting for the c-myc tag, which showed a band at the same size as that of the purified protein. Mass spectrometry analysis of the purified proteins also confirmed the identity of these two protein domains, providing 14 unique peptides for EDC (27% coverage) and 31 peptides for EGF (77% coverage).

**Figure 3:**
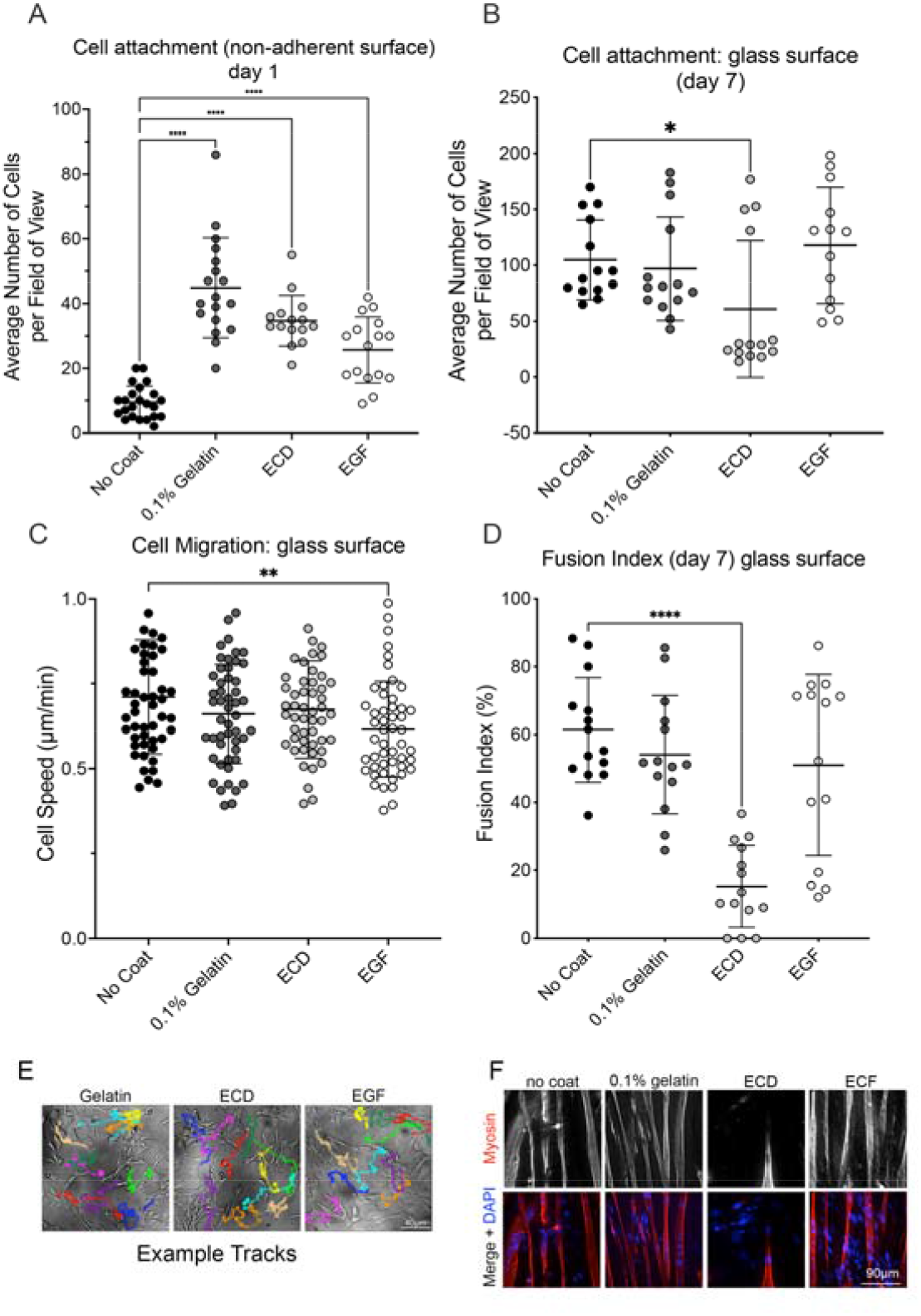
Effects of purified extracellular MEGF10 domains on myoblast fusion attachment and migration. **A:** Comparison of the ability of no-coat, 0.1% gelatin coating and purified ECD and EGF domains from MEGF10 to promote C1F myoblast attachment to non-adherent growth surfaces. Attachment was measured after 24 hours. **B**: Comparison of C1F myoblasts attachment to glass surfaces not coated, or coated with 0.1% gelatin, purified ECD or EGF domains after 7 days of differentiation. Individual measurements in **A** and **B** represent the numbers of cells per field of view (minimum of 5) collated from 3 biological replicates. **C:** shows the migration (cell speed) of myoblasts on these different coated surfaces 24 hours after plating (individual tracks from 3 biological replicates) and **D:** shows fusion on these differently coated glass surfaces at day 7 of differentiation. All the data show multiple individual measurements for three biological replicates. The mean ± S.D. of the mean is shown together with the results from an ANNOVA. * *P*<0.05 ** *P*<0.01, **** *P*<0.001 **E**. shows example images for myoblast motility/tracking and **F** shows example images for myoblast fusion on the different surfaces at day 7.

**Figure 4.**
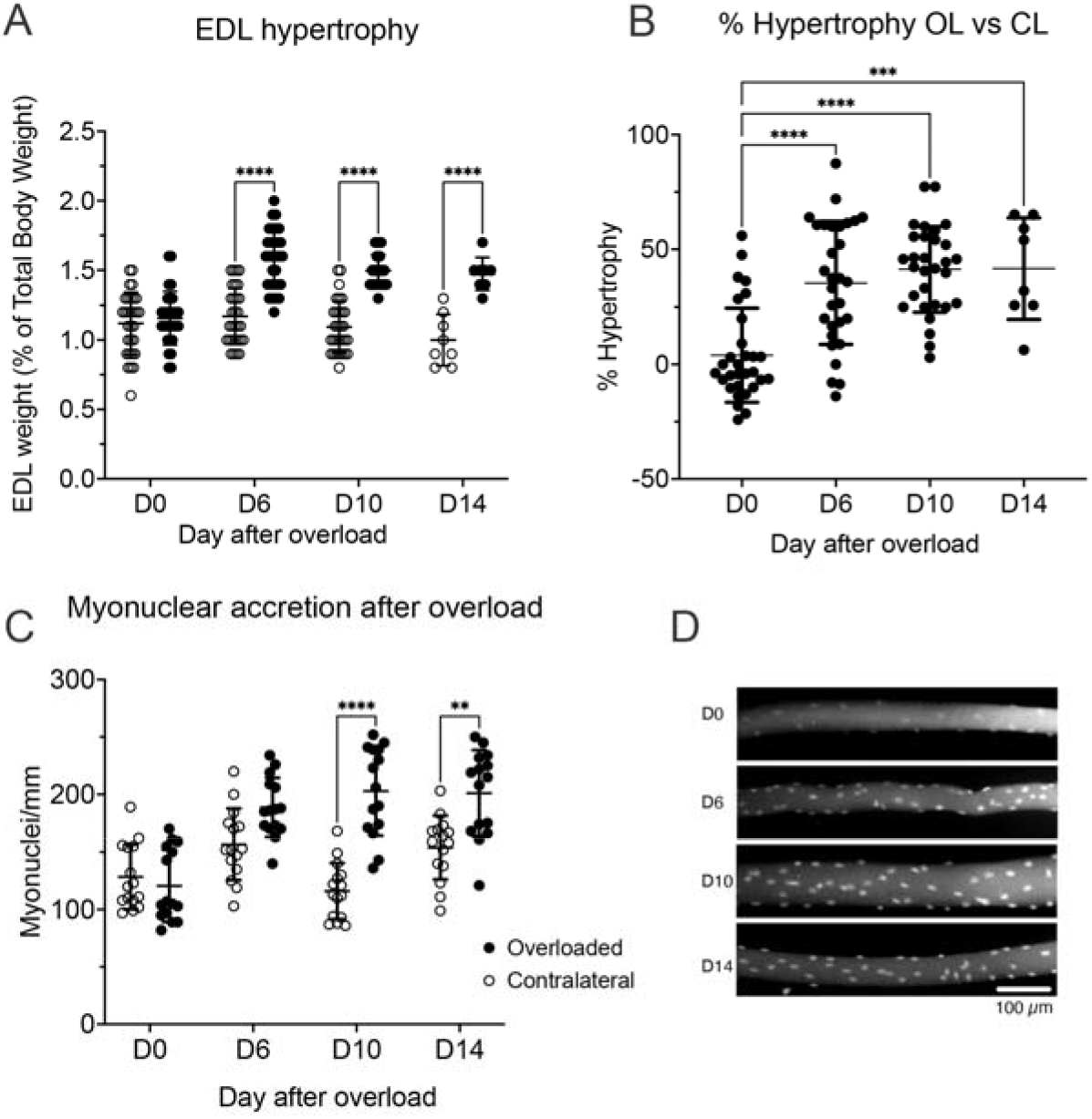
Variation in EDL hypertrophy and number of myonuclei following overload as a function of time in wild type mice. **A**. The relative weight (EDL muscle weight as a proportion of the total mouse body weight) is shown (mean values +/-S.D.) at each time point for unloaded contralateral (CL) and overloaded (OL) EDL (n=30 for D0, D6 and D10; n=6 for D14). **B**. Hypertrophy expressed as the % change in weight between unloaded CL and OL EDL. **C**. The numbers of myonuclei per 1mm of fibre from overloaded and contralateral EDL muscle at different days after overload (N=15 fibres, from 3 biological replicates). All the data was analysed using a 2way ANOVA (Sidák’s multiple comparisons test) **** *P*<0.0001. *** *P* <0.001. ** *P* <0.01. **D**. Representative mages of overloaded muscle fibres after different days of overload. Nuclei stained with DAPI.

The increased molecular weights of the expressed and purified EGF and ECD domains are likely to be the result of post-translational modifications such as glycosylation. Mass spectrometry showed that the ECD contained O-GlcNac modified residues. Using lectin blots (Supplemental Fig. 3C), we found that the expressed ECD and EGF were both highly glycosylated. This both explains the increased molecular weight compared to that expected and demonstrates that these domains are likely to be post-translationally modified in a similar way to what would be expected for endogenous MEGF10.

Next, we used these purified domains to coat non-adherent culture plates, and tested if they could promote cell attachment, using gelatin as a control. We found that both ECD and EGF domain constructs increased cell adhesion to non-adherent plates, compared to no coating at all (Fig. 3A), suggesting that both domains can promote cell adhesion of myoblasts.

We then coated glass surfaces with gelatin, ECD and EGF constructs to test if these domains promoted cell adhesion on adherent surfaces and had effects on differentiation as observed for exogenous eGFP-MEGF10 expression. At day 7 of differentiation, the total number of cells per area was significantly reduced for the ECD construct compared to cells differentiated on uncoated surfaces, and the fusion index was also markedly reduced (Fig 3B, D) but there were no significant effects on cell migration (Fig. 3C). Thus, addition of the external ECD domain has somewhat similar effects to expression of eGFP-MEGF10. However, the EGF construct (which lacks the EMI domain) did not affect number of cells at day 7 of differentiation or fusion, but did reduce cell migration (Fig 3B-D).

Taken together, these data suggest that MEGF10 expression must be tightly regulated for its correct cellular function and that the EMI domain is more important in modulating cell adhesion and fusion than the EGF domains. The EMI domain appears to be important in promoting adhesion of cells to a non-adhesive surface, but likely inhibits cell-cell adhesion required for fusion, thus reducing cell number at day 7 of differentiation and reducing fusion.

### Effect of muscle overload on the EDL muscle in wild type, *Megf10* ^+/-^ and *Megf10* ^-/-^ mice

We next performed experiments using genotyped wild type, *Megf10* ^+/-^ and *Megf10* ^-/-^ mice to understand the potential *in vivo* roles of MEGF10. The phenotype of MEGF10 knockout mouse has been reported to be relatively mild and exacerbated when crossed to the dystrophin knockout mouse model (mdx) (Saha et al., 2017). The same study showed that injection of barium chloride into the tibialis anterior muscle, an approach that results in severe muscle damage (Hardy et al., 2016), reduced the rate of fibre regeneration in MEGF10^-/-^ mice. Here, we focussed on the response of skeletal muscle to stimuli that elicit hypertrophy, a milder treatment. In this case, the TA muscle is removed, to induce overload on the EDL (extensor digitorum longus) muscle, which hypertrophies as a response. This can occur both as an increase in protein synthesis and though stimulation of satellite cells (Murach et al., 2021). In all of these experiments, the tibialis anterior muscle is only removed from one leg, resulting in overload (OL) on the EDL muscle in this leg. In the contralateral (CL) muscle, the TA muscle is kept intact, acting as a control.

Test experiments revealed that the overload response peaked at 10 days after surgery (Fig. 4: D10). The relative mass of the overloaded EDL muscles increased at each of the three time points sampled: D6, 10 and 14 compared to D0 (Fig. 4A). In contrast, the relative mass of the unloaded contralateral EDL muscles did not change, with no significant difference between contralateral tissue at all time points compared to D0 EDL (Fig. 4A). Expressing the change in weight as % hypertrophy comparing the overloaded muscle (OL) to the contralateral muscle (CL) (Fig. 4B) shows that the overloaded muscles hypertrophied by ∼40% at D6-14, a significant increase compared to D0. Muscle hypertrophy appeared to plateau between days 10 and day 14.

Counting the numbers of nuclei per fields of view for isolated muscle fibres showed that the numbers of nuclei per mm was significantly increased in the overloaded compared to contralateral muscle fibres at D10 and D14. The percentage increase in nuclei per mm in overloaded fibres was significantly increased compared to D0 (Fig. 4C, D). No significant differences were found when number of nuclei on contralateral fibres were compared between time points, or to D0 fibres. The increase in numbers of nuclei shows a similar trend to the increase in EDL mass following overload. As the largest effects of overload were observed at D10, this time point was used in subsequent experiments.

### The hypertrophy response to overload is reduced in Megf10^-/-^ mice

After 10 days, the weight of the overloaded EDL muscle from wild type and *Megf10*^+/-^ mice increased significantly compared to the contralateral, unloaded muscles but was not significantly different for homozygote *Megf10*^-/-^ mice (Fig. 5A). However, the percentage change in hypertrophy was lower for overloaded EDL muscles from *Megf10*^+/-^ and *Megf10*^-/-^ mice compared to wild type (Fig. 5B). Similarly, there was a significant increase in myonuclear number (myonuclear accretion) in overloaded EDL muscles from all three genotypes, which was similar in magnitude for wild type and *MEGF10*^+/-^ mice. However, myonuclear accretion was significantly reduced in overloaded EDL muscles from *Megf10*^-/-^ mice (Fig. 5C). These results suggest the complete loss of MEGF10 reduces the hypertrophic response to muscle overload. The overall fibre cross sectional area (CSA) per mouse did not significantly change, although there was a trend for fibre CSA to increase for wild type and *MEGF10*^+/-^ mice, but not for the *Megf10*^-/-^ mouse.

**Figure 5.**
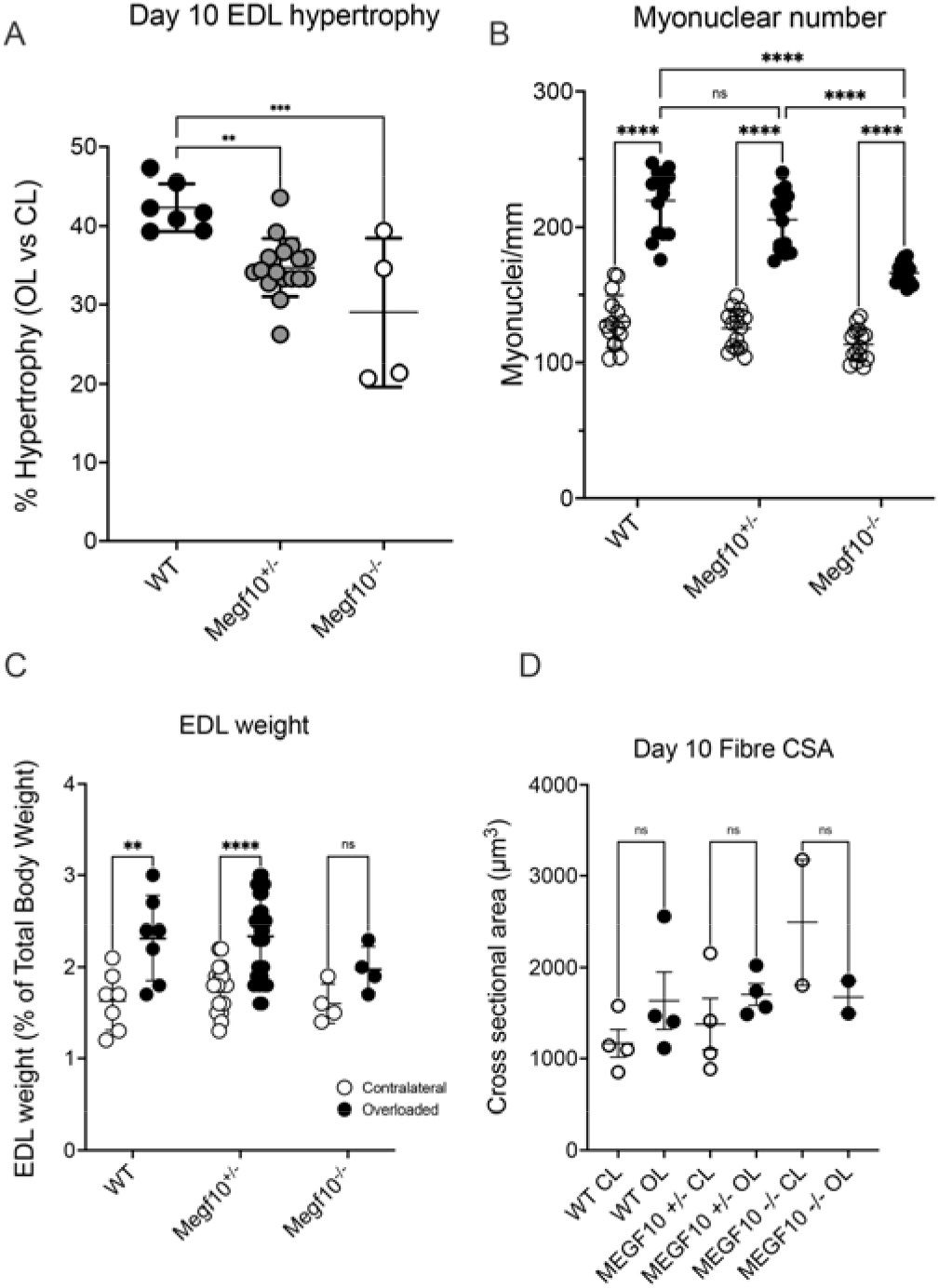
*Megf10*^+/-^ and *Megf10*^-/-^ mice show reduced EDL hypertrophy following muscle overload: day 10. **A**. The relative weights for unloaded (contralateral) and overloaded EDL muscles for wild type, heterozygous and homozygous knockout Megf10 mice. B: Relative hypertrophy (%) of the overloaded EDL muscle following 10 days of overload. C. numbers of myonuclei per 1mm of fibre from overloaded and unloaded (contralateral) EDL muscles at day 10. (n=15 fibres, minimum of 3 biological replicates) D. Mean fibre cross sectional area of muscle fibres (per mouse). Data was analysed by ANOVA. Error bars represent S.D. ** *P* <0.01, **** *P*<0.001, **** *P* <0.0001. ns: non-significant.

### Lower numbers of Pax7^+^ cell nuclei are present at D0 and increases in transcription factor expression induced by overload are reduced in in Megf10^+/-^ and Megf10^-/-^ mice

Loss of MEGF10 expression in humans has been linked to decreased numbers of Pax7^+^ satellite cells (Logan et al., 2011). Decreased numbers of Pax7^+^ satellite cells could underlie the reduced hypertrophic response observed for *Megf10*^-/-^ EDL muscle. To evaluate this, fibres were isolated from mice and stained for the transcription factors Pax7, MyoD and myogenin, at D0 and after 10 days of overload. At day 0, the numbers of Pax7^+^ positive satellite cells were lower in heterozygote *MEGF10*^+/-^ and homozygous knockout *MEGF10*^-/-^ mice compared to wild type (Fig. 6A). A similar, but non-significant trend was found for MyoD and myogenin (Fig. 6A). This suggests that the numbers of satellite cells are reduced in EDL from *MEGF10*^+/-^ and *MEGF10*^-/-^ mice at D0.

**Figure 6.**
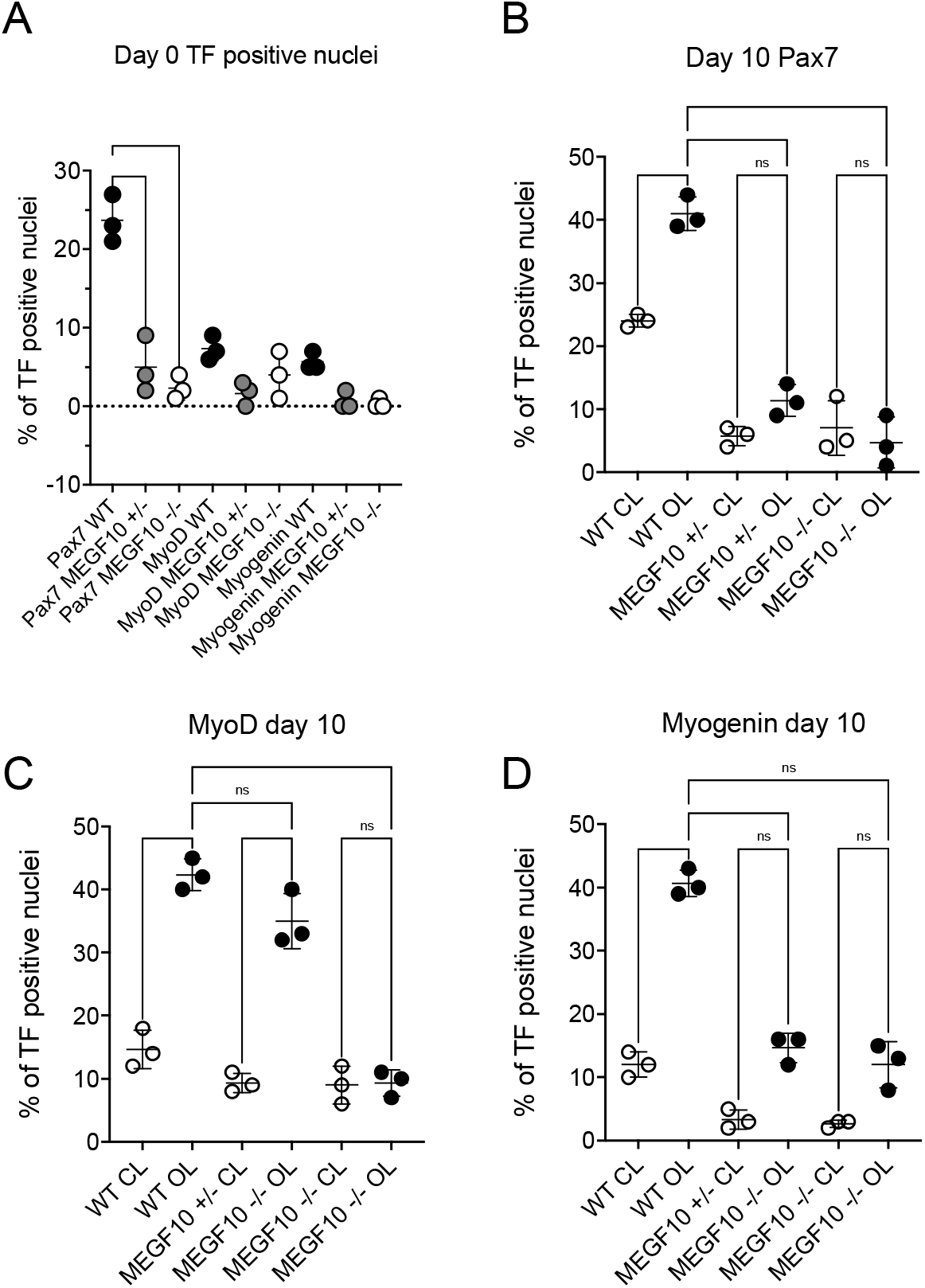
Percentage Transcription Factor (TF) expression by satellite cells on fibres from wild type, Megf10^+/-^ and Megf10^-/-^ mice. A: Expression of each TF (Pax7, MyoD and Myogenin) at day 0 for wild type, MEGF10^+/-^ and MEGF10 ^-/-^ mice. **B-D** shows the expression of each TF for contralateral, unloaded and overloaded EDL muscles at day 10. Values shown in **A-D** are the mean values per mouse (using the EDL muscle) together with the S.D. Data was analysed by ANOVA. **** *P* <0.0001. * *P* <0.05. ns: non-significant.

After 10 days of overload, the percentage of nuclei that stained positive for Pax7, MyoD and myogenin increased significantly for overloaded EDL muscle from wild type mice compared to the contralateral (CL) muscle (Fig. 6B-D). However, the percentage of nuclei positive for Pax7 and myogenin did not increase significantly in overloaded EDL muscle from *MEGF10*^+/-^ and *MEGF10*^-/-^ mice compared to the unloaded, contralateral muscle (Fig. 6B, D). The number of MyoD positive nuclei for the overloaded EDL muscle did increase significantly for *Megf10*^+/-^ mice but not for *Megf10*^-/-^ mice (Fig. 6C). In summary, overload of the EDL muscle in *Megf10*^+/-^ and *Megf10*^-/-^ mice reduces the increase in transcription factor expression compared to wild type mice.

### Mean fibre cross-sectional area in the diaphragm from Megf10^+/-^ mice is smaller

As we did not obtain the expected Mendelian ratio for the genotypes of pups at birth, and to further identify effects of the loss of Megf10 on muscle, we analysed the diaphragm muscle from 6-week old wild type and *Megf10*^+/-^ mice. The muscle was stained for laminin to outline the muscle fibres and show if muscle fibre size and organisation is affected. This revealed strong differences between wild type and *Megf10*^+/-^ mice (Fig. 7A). The mean fibre CSA of individual fibres from the diaphragm muscles were reduced significantly in *Megf10*^+/-^ compared to wild type mice (Fig. 7B). The thickness of the laminin in the extracellular matrix between fibres was increased significantly in the *Megf10*^*+/-*^ diaphragm compared to wild type (Fig. 7D). This may explain why the Mendelian ratio was not as expected, as it is possible that the diaphragm of *Megf10*^*-/-*^ mice is more strongly affected, and mice do not survive long after birth.

**Figure 7.**
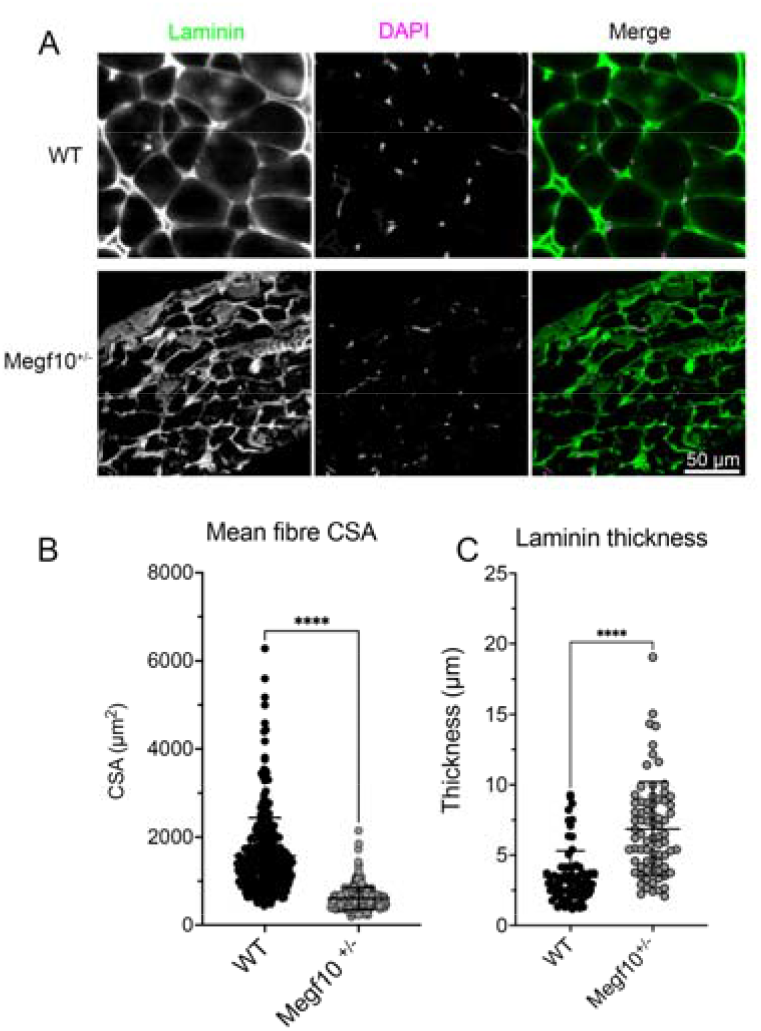
Comparison of wild type and Megf10^+/-^ diaphragm. **A**. Representative images of stained diaphragm cross-sections showing differences in fibre structure. B. Scatter plot showing individual measurements of CSA across 3 biological replicates. C. Scatter plot showing thickness of laminin divisions between fibres, measured from 5 fibres per mouse. Bars show mean ± S.D. Error bars represent S.D. **** *P* <0.0001. *** *P* <0.001. * *P* <0.05.

## Discussion

Here, using a combination of *in vitro* and *in vivo* experiments, we have shown that overexpression of GFP-MEGF10 or addition of the extracellular domain of MEGF10 (ECD) inhibits fusion of C1F myoblasts. The purified full-length ECD was more effective in attachment to a non-adhesive surface than the shorter EGF domain, in which the N-terminal EMI domain was absent. However, it was less effective at maintaining cell-cell attachment as cell differentiate into myotubes, reducing both number of cells and fusion at day 7 of differentiation, suggesting a key role for the EMI domain in these processes. *In vivo*, loss of Megf10 reduced hypertrophy of the EDL muscle in response to overload, as measured by weight change and myonuclear accretion, with the most marked effect for homozygous knockout *Megf10*^*-/-*^ mice. This reduced response may be accounted for by the reduction in Pax7^+^ satellite cells, and decreased satellite cell response to overload in Megf10 heterozygous (*Megf10*^*+/-*^) and homozygous knockout mice. Finally, homozygous knockout mice were born at a lower mendelian ratio than expected. An analysis of the diaphragm muscle in wild type and heterozygous mice showed more variable and reduced fibre cross-sectional area in the heterozygous mice compared to wild type, and increased laminin deposition between fibres. The lower survival rate in homozygous mice, could arise from increased defects in the diaphragm muscle, which reiterates the main clinical presentation of affected individuals with MEGF10-related EMARDD; respiratory distress due to diaphragmatic paralysis.

During myoblast differentiation into myotubes, previous work demonstrated that that *Megf10* is downregulated during differentiation of C2C12 cells, by qPCR (Holterman et al., 2007) and in C1F myoblasts differentiated on glass surfaces, by RNAseq analysis (Richardson et al., 2022). These changes have not been confirmed at the protein level, as a well-validated antibody to Megf10/MEGF10 is lacking. However, if Megf10 is normally downregulated during myoblast differentiation, then overexpression of MEGF10 *in vitro*, either *via* expression of eGFP-MEGF10 or via addition of the purified ECD of MEGF10 might be expected to interfere with fusion and differentiation, as we found here. Our findings for eGFP-MEGF10 are similar to those observed previously for overexpression of HA-tagged Megf10 in C2C12 myoblasts, which also decreased myoblast fusion and differentiation *in vitro* (Holterman et al., 2007).

Addition of purified ECD domain also inhibited fusion of myoblasts in vitro and was more effective at inhibiting fusion than the EGF domain, which lacks the EMI domain. However, the ECD domain was more effective at promoting cell attachment to non-adhesive surfaces than the EGF domain. Thus, the EMI domain seems to be important in both inhibiting fusion (which requires cell-cell adhesion) and in promoting cell-surface adhesion. The EMI domain is found in many other MEGF isoforms, and also in the worm orthologue of human MEGF10 (Callebaut et al., 2003). If the EMI domain is able to recognise and bind to PS, then its presence in fusing cells that expose PS on their surface could potentially block binding of other PS receptors that promote fusion (Chikazawa et al., 2020; Hochreiter-Hufford et al., 2013; Jeong and Conboy, 2011; van den Eijnde et al., 2001). This suggests a possible role of MEGF10 in preventing premature myoblast fusion. Of note, MEGF10 has also been suggested to interact with Notch *via* its intracellular domain (reviewed in (Vargas-Franco et al., 2022)), and thus the effects of the exogenous ECD and EGF domains would be independent of any downstream signalling pathway involving Notch.

Our observation of a hypertrophic response of the EDL muscle to overload for wild type mice is consistent with previous reports (Egginton et al., 2011; Johnson and Klueber, 1991). The increase in mass of the overloaded EDL as well as the increase in myonuclei has been reported previously (Bruusgaard et al., 2010; Huey et al., 2016; Seiden, 1976; Zhou et al., 1998). Moreover, the activation of satellite cells by overload, with increases in Pax7^+^, MyoD^+^ and myogenin^+^ satellite cells of wild type mice matches findings from previous reports (Hyatt et al., 2008; Sakuma et al., 1999). However, we did not find a strong effect of overload on fibre cross-sectional area, although this has been observed previously in other overload conditions (Carter et al., 1995; Rosenblatt et al., 1994; Snijders et al., 2020). Overall, our experimental data shows that EDL from wild type mice responds to physiological overload as expected, and that this is a suitable approach to explore the effects of overload in *Megf10*^+/-^ and *Megf10*^-/-^ mice.

The reduction in or loss of Megf10 reduced the hypertrophic response of the EDL muscle to overload for both heterozygous *Megf10*^+/-^ and homozygous knockout *Megf10*^-/-^ mice, compared to wild type mice. This reduction was greatest for the homozygous mice. Moreover, the increase in Pax7, MyoD and myogenin positive cells was reduced, again with the largest effect in homozygous mice. The contribution of satellite cells to hypertrophy has been the subject of much discussion (Pallafacchina et al., 2013) and it has been shown that some increase in fibre size is possible without functional satellite cells (Murach et al., 2017). However, Pax7 expression has been reported to contribute to the activation and subsequent expansion of satellite cells in response to stimuli such as overload (Wang and Rudnicki, 2011). Moreover, our findings agree with a previous report that found Megf10 deficient mice to have reduced expression of Pax7 and MyoD, resulting in inadequate regeneration of EDL muscle following acute injury due to barium chloride treatment (Li et al., 2021).

In addition to the reduced hypertrophic response, we also found that muscle fibres in the diaphragm were highly variable in size, and that there was increased fibrosis. Loss of MEGF10 in humans causes respiratory distress (Logan et al., 2011). The change to the structure in the diaphragm of Megf10^+/-^ mice we observed could also lead to respiratory distress and could help to account for the lower numbers of homozygous mice that we obtained. Further work is needed to confirm if the diaphragm of Megf10^-/-^ mice is affected more severely. Interestingly, a recent report described defects in the neuromuscular junction (NMJ) in the tibialis anterior and diaphragm muscles in Megf10^+/-^ mice, consistent with its role in glial cells in synapse remodelling (Juros et al., 2024), suggesting altered NMJs may also contribute to the phenotype of the diaphragm muscle seen here. However, the NMJ of the EDL muscle was not affected in Megf10^-/-^ mice (Juros et al., 2024), suggesting that alterations to the NMJ do not have a major contribution to the reduced hypertrophic response we observed here.

Overall, our results demonstrate that MEGF10 is likely to be important in myoblast-surface adhesion but can potentially block fusion of myoblasts when present at high levels, thus possibly preventing premature fusion of satellite cells. The reduction in numbers of Pax7^+^ satellite cells, and the reduction in their activation in response to muscle overload in *Megf10*^*-/-*^ mice broadly supports this idea. Satellite cells could be lost through poor adhesion and/or premature fusion. The overall phenotype of the *Megf10*^*-/-*^ mice recapitulates the human phenotype observed in EMARDD (Logan et al., 2011), resulting from homozygous nonsense mutations in the *MEGF10* gene. In addition to respiratory distress resulting from paralysis of the diaphragm muscle, the reduced numbers of Pax7^+^ satellite cells, reduced skeletal muscle fibre growth and reduced satellite cell activation in the mouse model, are reminiscent of the reduced number of Pax7^+^ satellite cells, small muscle fibres, fibre necrosis and fibre replacement by fibrous or adipose tissue in humans (Logan et al., 2011).

## Materials and Methods

### *In vitro* experiments: cell culture and staining

To explore muscle differentiation *in vitro*, we used a clone of satellite cells (C1F) derived from the *H-2k*^*b*^-tsA58 mouse (Morgan et al., 1994; Richardson et al., 2022). These cells proliferate at 33°C in the presence of IFN□ in growth medium (Dulbecco’s minimal essential medium (DMEM), high glucose, containing Glutamax (Gibco), supplemented with 20% FCS, 1% Penicillin/Streptomycin (v/v) (Gibco), 2% Chick Embryo Extract (E.G.G. Technologies)) and are switched to differentiate by changing the medium to DMEM supplemented with 4% Horse Serum and 1% penicillin/streptomycin, and increasing the temperature to 37°C. Proliferation ceases within 24 hours, and cells start to fuse at this time point. RNASeq experiments suggested that Megf10 is expressed in these cells (Richardson et al., 2022).

For immunostaining, cells were plated onto washed 13 mm diameter glass coverslips, coated with 0.1% gelatin. Cells were fixed with 4% paraformaldehyde (PFA) in phosphate buffered saline (PBS) for 20 minutes, washed in PBS, permeabilised with 0.5% Triton X-100 diluted in PBS containing 1% (w/v) bovine serum albumin (BSA) for 30 minutes prior to incubating with primary antibodies (Supplementary Table 1) diluted into PBS with 1% (w/v) BSA for 60 minutes at room temperature. Coverslips were washed x5 with PBS, and then appropriate anti-mouse or anti-rabbit secondary antibodies (Alex-Fluor conjugated, Invitrogen) diluted 1/400 in PBS with 1% (w/v) BSA were added. Fluorescent phalloidin was used to stain filamentous actin and DAPI (4′,6-diamidino-2-phenylindole) was used to stain nuclei (Sigma). Of note, we were unable to validate any commercial antibody to MEGF10, and thus could not stain or blot for endogenous protein. Once stained, coverslips were mounted onto glass microscope slides using Prolong gold antifade mountant (Invitrogen). Cells were imaged using a Zeiss LSM Airyscan confocal microscope, using a x40 objective lens (NA 1.4) or an Olympus widefield microscope using a x63 objective lens (N.A. 1.4) followed by deconvolution. The resulting images were assembled into figures using Adobe photoshop.

### GFP-MEGF10 adenoviral construct generation

Adenoviral expression constructs for GFP-MEGF10 (full length human MEGF10, 3423 bp was provided by Colin Johnson) and for GFP were generated by PCR-based cloning into a pDC315 vector (Addgene). For GFP-MEGF10, eGFP was placed at the C-terminus of MEGF10, separated from MEGF10 by a short 3x glycine linker, and a 6xHis tag was incorporated at the C-terminus after the eGFP (Fig. 1B). Fully sequenced constructs, with no sequence errors, were used to generate adenovirus using Ad293 cells (Microbix). Purified DNA plasmids (pDC315 alongside pBHGloxΔE1,3Cre) were transfected into Ad293 cells, using Fugene and the resulting adenovirus was amplified as described in the manufacturer’s protocol (Wolny et al., 2013). After the final round of amplification, virus was purified using the Vivapure Adenopack 100 kit (Sartorius Stedim Biotech), purified virus was stored in storage buffer (20 mM Tris/HCl, 25 mM NaCl, 2.5% glycerol (w/v), pH8) at -80°C. Viral titre was determined by tissue culture infectious dose 50 (TCID50) method. Final titres for GFP-MEGF10 and the GFP virus were 3 x 10^8^ and 6 x 10^8^ PFU per ml respectively. This was used to estimate the MOI of infection, when infecting cultured myoblasts. Western blots confirmed the expected sizes for GFP-MEGF10 and GFP (Supplemental Fig. 1).

### Expression and purification of extracellular domain constructs of MEGF10

Extracellular MEGF10 constructs (human) were generated for mammalian protein expression (Fig. 1B). The cDNA for the extracellular domain (ECD: residues Leu35 to Gly861) or the ECD lacking the EMI domain (ECF, His 108 to Gly861) were cloned into the pSecTag2A plasmid (Invitrogen), sequences confirmed, and constructs were transfected into HEK-293 cells using calcium chloride transfections. pSecTag2a contains an N-terminal secretion signal from Ig-κ for efficient protein secretion in the media, a cytomegalovirus (CMV) promoter for high-level expression and C-terminal 6xHis and c-myc tags for nickel column purification and antibody detection, respectively, as well as a Zeocin resistance gene for the selection of stably expressing mammalian cell lines. 48 hours after transfection, cells were harvested, diluted to 1 x 10^5^ cells per ml, and seeded at 2 x 10^4^ cell per ml in selection medium (DMEM, 10% FCS, 1 % FCS, 200 μg/ml Zeocin). After 10 days, individual clones, were picked into 24 well plates and allowed to grow. Samples of media from each colony were analysed by dot blot, to isolate clones for which expression was highest.

For expression and purification of the MEGF10 extracellular domain constructs, stable cell lines with high levels of expression, were seeded into five 75 cm^3^ flasks coated with 20 µg ml^-1^ poly-L-lysine, grown to 80% confluence in normal growth medium and then the medium was exchanged for OptiMEM low serum medium (GIBCO) to reduce contaminants during purification, and cells cultured for a further 3 days. The medium was then removed, centrifuged at 1000 x *g* rcf, and the supernatant incubated with 1 ml Complete His-Tag Purification Resin slurry (Roche) and Complete EDTA-free protease inhibitor cocktail tablet (Roche) for 30 min on a roller. The mixture was then applied to a 5 ml column, and the flow through collected. The resin was washed 5x with column wash buffer (300 mM NaCl, 50 mM NaHPO_4_) and eluted with elution buffer (300 mM NaCl, 50 mM NaHPO_4_, 200 mM Imidazole). Eluted protein (in 2 mL) was dialysed into PBS overnight using a Gebaflex Maxi Dialysis Tube (MWCO = 3.5kDa) (Generon). Purified protein was stored in 200 µL aliquots at -80 °C. Protein concentration was measured using a Cary 50 Bio-UV visible spectrophotometer (Varian) at a wavelength of 280 nm. Typical concentrations were 60-100 ng/µl for ECD and 150-200 ng/µl for ECF. Protein identity was confirmed by mass spectrometry. A lectin blot (Biotinylated Lectin Kit I (Vector Laboratories)) was used to confirm that the ECD and EGF constructs had been glycosylated.

### Analysis of protein expression

Samples of cells used for western blotting were either prepared by scraping cells directly into 2x Laemlli buffer prior to boiling at 100 °C for 10 min and freezing aliquots at -80 °C, or freshly pelleted cells were resuspended into 50 µl ice-cold lysis buffer (150 mM NaCl, 50 mM Tris (pH8), 1% Triton X-100, 1 mM EDTA (pH8)) containing Halt Protease Inhibitor, single-use cocktail (ThermoScientific), incubated on ice for 30 mins with regular vortexing, centrifuged at 17000 x *g* rcf for 20 mins at 4 °C, and lysates stored at -20 °C. Protein concentration was then quantified by a Pierce micro BCA Protein Assay kit (Thermo Scientific) following the manufacturer’s instructions. Absorbance at a wavelength of 544nm was measured using a Polstar Optima plate reader.

### Analysis of myoblast fusion and cell motility

To analyse cell motility, cells were imaged for 14 hours, capturing images every 10 minutes, using differential interference contrast (DIC) microscopy (Olympus widefield microscope), x10 objective lens at 37 °C. For eGFP-MEGF10 expressing cells, individual wells of a 96 well plate with a borosilicate glass bottom (Iwaki) were seeded with 50 μl of C1F cells at 1×10^5^ cells/ml and infected overnight using an MOI of 100 for each adenoviral construct in 500 μl culture media. For each condition, 5 fields of view were imaged and the experiment was repeated three times (3 biological replicates). Cell motility was analysed using ImageJ software (MTrackJ plugin), tracking 10 cells per field of view. To determine the effect of the purified ECD and ECF domains on cell motility, individual wells of the borosilicate glass were coated with 1.25 μg of purified protein for 20 min at 37°C. Excess coating was aspirated and wells seeded with 50 μl C1F cells at 1×10^5^ cells/ml. Cells were incubated with 50 μl of culture media at 33 °C, 10% CO_2_ for 24 hrs, medium made up to 500 μl and then cells were filmed and motility analysed as above.

To estimate the fusion index, C1F cells were seeded onto pre-washed 13 mm diameter coverslips, coated with 0.1% gelatin and differentiated for seven days. Cells were fixed with pre-warmed 4% PFA for 20 minutes, washed with PBS and then permeabilised with 0.1% Triton X-100 in PBS. The nuclei were stained for nuclei using DAPI, filamentous actin using fluorescently labelled phalloidin and striated muscle myosin using the A4.1025 antibody (Cho et al., 1994; Maggs et al., 2000). The fusion index is calculated from the percentage of nuclei found in skeletal myosin positive myotubes as a percentage of the total number of nuclei. Only skeletal myosin positive myotubes with three or more nuclei were classed as myotubes. For each condition, five fields of view were imaged and cells counted from three biological replicates.

### Cell attachment and cell motility assays

To determine the ability of the ECD and ECF proteins to mediate C1F myoblast attachment to a surface a cell attachment assay was performed. Briefly, the wells of a non-adhesive 24 well plate (Greiner), were coated with 2.5 μg protein diluted in 150 μl PBS for 20 min at 37 °C. Harvested myoblast cells (C1F clone) were prepared at 1×10^4^ cells ml^-1^ in growth medium and 100 μl added to each well. Cells were incubated at 33 °C, 10% CO_2_ for 30 min before adding 500 μl of fresh medium. After 24 hr incubation, cells were imaged using a Cytomate instrument, taking 5 different fields of view and counting the number of nuclei from each field. The experiment was repeated three times. Alternatively, a 96 well borosilicate glass plate (Iwaki) was coated for 20 minutes at 37 °C with 0.1% gelatin, or with 1.25 µg of the EGF or the ECD domains of MEGF10, or left uncoated, coating was aspirated and wells seeded with 50 µl of CIF cells at 1 x 10^5^ cells per ml. Cells were allowed to attach, supplemented with additional medium (50 µl), and filmed overnight to analyse their motility.

### Megf10 knockout mice

Megf10tm1(KOMP)Vlcg mice (RRID: MMRRC_048576-UCD, MGI ID: 4454190, background: C57BL/6Tac, intragenic targeted knockout deletion, gene ID: 70417) were obtained from the Mary Lyon Centre at the MRC Harwell Institute and exported to our laboratory to establish breeding colonies. Briefly, the model was generated by replacing exons 1-24 of mouse *Megf10* by homologous recombination with an expression selection cassette as detailed by the Knockout Mouse Project (KOMP, University of California Davis, Davis, CA: https://www.komp.org/geneinfo.php?geneid=68051). The *Megf10*^tm1(KOMP)Vlcg^ mice harbour the Velocigene cassette ZEN-Ub1 inserted into the *Megf10* gene between positions 57340143 and 57372060 on chromosome 18, generating a 31918bp deletion that deletes exons 1-24 of MEGF10. The mouse line was rederived in the Harwell facility after its original generation at Regeneron Pharmaceuticals. qPCR has shown no mRNA for MEGF10 is expressed in the cerebellum in homozygote animals (Mouse Genomics Informatics) (Iram et al., 2016) and knockouts, hereafter designated *Megf10*^*-/-*^, lack MEGF10 protein (Fig. 1f in (Chung et al., 2013)), confirming that the *tm1* allele is null. *Megf10*^*-/-*^ (homozygous knockout) mice were generated on a C57BL/6NTac background. *Megf10*^*-/-*^, *Megf10*^*+/-*^ (heterozygous), and wild type animals used in the experiments were generated by crossing *Megf10*^*+/-*^ heterozygotes, and all comparisons between genotypes are between age-matched littermates. Male and female adult C57BL/6Tac (bred in-house at the University of Leeds) and *Megf10*^tm1(KOMP)Vlcg^ mice (final body mass approx. 25g) were used in this study.

All experimental procedures and sacrifice were conducted with approval of the local animal welfare and ethics committee, under Home Office project licences 70/8674 and PP1775021. Mice were housed in groups in a temperature-controlled environment with access to food and water *ad libitum*. Cages underwent 12/12 light/dark cycles. Breeding was carried out under service licence PP0237211, with two breeding cages of C57BL/6Tac x heterozygous *Megf10*^tm1(KOMP)Vlcg^, and one breeding cage of heterozygous *Megf10*^tm1(KOMP)Vlcg^ x heterozygous *Megf10*^tm1(KOMP)Vlcg^. Mice were stunned by concussion and killed by cervical dislocation.

### Genotyping

To accurately determine the genotype for each mouse, ear biopsies were collected by unit staff at Central Biomedical Services (CBS), University of Leeds. Biopsies were placed into 96-well plates, sealed, and shipped to Transnetyx for genotyping (Transnetyx Inc. Cordova, TN) *via* courier. A bespoke PCR assay to determine genotype was designed by the Genetic Services team at Transnetyx based on information provided by KOMP (Knockout Mouse Project) mouse repository (Supplementary Fig. 1). Results were obtained within 72 hours of sending the samples. Of note, we did not obtain the expected Mendelian ratio of 1:2:1 (Supplemental Table 2).

### Hypertrophy model surgery

Unilateral extirpation (removal) of tibialis anterior (TA) muscle was performed under aseptic conditions and inhalation anaesthesia. All instruments were sterilised and work carried out under a dissection microscope. Mice were first anaesthetised with 5% isoflurane in 2Lmin^-1^ O_2_. The left leg was then shaved and wiped with ethanol to sterilize the area and remove surface bacteria. For the remainder of the operation, mice were maintained under anaesthetic at 2% isoflurane in 2L/min O_2_. All possible steps were taken to avoid animals suffering at each stage of the experiment.

A single incision was made on the hindleg to expose the TA, tweezers used to lift the superficial distal tendon, and the TA removed by making incisions at proximal and distal points of attachment using a scalpel. The TA cut end was then held over the wound area for approx. 10s to allow the release of chemokines to aid repair and blood clotting. The TA was then discarded and 1-2 drops of 2.5% Baytril (Bayer AG) was applied to the wound for antiseptic protection. The area was swabbed with a cotton bud to remove blood and then the incision was sutured with MERSILK^TM^ (Ethicon Inc.) braided silk suture, size 5.0. Sutures were intermittent and double-knotted to reduce the chance of mice unravelling them post-operatively. 1-2 more drops of Baytril⍰ were applied to the closed wound and swabbed with sterile cotton buds, to remove dried blood that may lead to irritation. 0.1ml 10% Vetergesic (Ceva Animal Health, Ltd) was administered to the scruff of the neck to provide post-operative analgesia. Mice were placed in a heated cage without sawdust for approximately 10 minutes to recover from anaesthetic, before being placed back into their original cage. Mice were observed to be normally ambulant, thus overloading the extensor digitorum longus (EDL), for a pre-determined length of time before sampling.

### EDL isolation

Changes in EDL muscle phenotype were assessed in control animals (no overload), as well as animals overloaded for 10 days, to observe changes in the muscle. This interval was chosen as we observed that the overload response had peaked at this time point in test experiments. Mice were killed by Schedule 1 (concussion followed by cervical dislocation). Muscle was removed as quickly as possible to minimise post-mortem biochemical changes. The leg of a freshly killed mouse was shaven and dabbed with ethanol to promote cutaneous vasoconstriction. A small incision from just lateral to the knee to the beginning of the hindfoot aided blunt dissection using scissors and forceps to break through the layer of fascia atop the muscle. On the unoperated (contralateral) leg, the TA was first removed to access the EDL underneath. The EDL was then accessed and removed in the same way (forceps used to hold the tendon and scalpel used to release it at the base). On the ipsilateral (overloaded) leg, the EDL was simply removed as described. The EDL, and the whole mouse, were both weighed to determine the EDL mass as a percentage of total body mass.

### Preparation of muscle samples for imaging

Samples of skeletal muscle were additionally harvested for single fibre isolation (as described above), or for cryo-sectioning. For accretion measurements, single fibres were fixed and permeabilised, incubated with DAPI (1/500) for 90 minutes at room temperature before washing with TBST and finally with PBS. Samples were mounted on cleaned glass microscope slides using ProLong Gold and covered by a 20 x 40 mm glass coverslip. For cryosectioning, intact muscles were trimmed, mounted onto a cork disk with optimum cutting temperature compound (OCT) (Agar Scientific) and immediately snap frozen in isopentane-liquid nitrogen and stored at a temperature of -80 °C. Diaphragm muscle was also prepared for cryo-sectioning, using a similar approach. Samples were sectioned (30 µm and 10 µm sections) using a cryostat (Leica) pre-cooled to -20 °C and sections placed onto labelled glass slides and stored at -20°C until ready to fix and stain.

Slides were left at room temperature for ∼10 mins to dissipate condensation. A hydrophobic barrier pen was then used to draw around each segment of 3-4 sections joined together. Within the confines of the hydrophobic barriers, tissue was fixed by applying 100-200 µl ice-cold 100% methanol and incubating at room temperature for 10 mins. Sections were then washed three times with PBS. Non-specific antibody binding was reduced by incubating with 5% BSA diluted in PBS at room temperature for 30 min. Primary antibodies were diluted in PBS and applied to tissue sections following removal of the blocking solution. Slides were incubated with the primary antibody overnight at 4°C. Primary antibody was removed and sections washed three times with wash buffer (1% BSA in PBS). Secondary antibodies were diluted in PBS, applied to tissue sections, and incubated for 1hr at room temperature. Secondary antibody was removed, and sections washed three times with wash buffer, before a final wash with PBS. Two drops of ProLong Gold antifade mountant was then added to the slide, and a 20 mm x 40 mm glass coverslip was placed on top. Slides were left overnight at room temperature in the dark overnight, and then stored at 4 °C.

### Quantification of transcription factor expression and fibre cross-sectional area

Stained fibres were imaged using a widefield Olympus IX-70 microscope, using a 40x, N.A. 1.4 objective lens. For measuring myonuclear accretion, 15 x DAPI stained fibres were imaged per condition (three fibres per EDL sample). Numbers of nuclei were counted from 1 mm sections using ImageJ (Fiji).

For transcription factor staining, the number of nuclei positively stained for a transcription factor (Pax7, MyoD or myogenin) per 50 myonuclei along the fibre was measured from three fibres per EDL sample, stained for DAPI and the transcription factor. The numbers of positive nuclei per fibre were expressed as a percentage of 50 myonuclei, and the number per fibre was averaged.

Cross-sectional area of individual muscle fibres was also determined using ImageJ (FIJI). Outlines of each individual fibre were drawn around on each image using the freehand selection tool, and area in µm^2^ was automatically calculated.

### Measurement of laminin thickness

The width of the basement membrane between fibres in stained diaphragm cross sections was measured using ImageJ processing software (NIH). The ‘straight line’ tool was used to take orthogonal measurements of laminin (visualised with anti-rabbit IgG Alexa Fluor 488 conjugate) normal to the sarcolemma, and this value recorded. Five measurements were taken per fibre, and five fibres were measured for 3 wild-type and 3 *Megf10*^+/-^ mice.

### Statistical analyses

Statistical tests were performed, and graphs generated, using GraphPad Prism for Mac (GraphPad Software, La Jolla California, USA, www.graphpad.com). Graphs show mean ± standard deviation (S.D.) for each observation. Unpaired t-tests with Welch’s correction and one-way and two-way ANOVAs were carried out to test for any statistically significant differences between conditions. The level of statistical significance is indicated by the number of asterisks displayed above graphs: **** represents a *P* value <0.0001. *** represents a *P* value <0.001. ** represents a *P* value <0.01. * represents a *P* value <0.05.

## Supplemental Material

The supplemental material comprises two supplementary tables: a table of the antibodies used in this work (Supplemental Table 1), a table of the Mendelian ratio (Supplemental Table 2) an outline of how mice were genotyped, and three supplemental figures: supplemental Fig. 1 the strategy used to genotype the mice, Supplemental Fig. 2: expression tests for eGFP and MEGF10-eGFP, and Supplemental Fig. 3: characterisation of the expressed and purified ECD and EGF domains, and their glycosylation.

### CReDiT

All authors contributed to the conceptualization of this work, methodology and writing, review and editing of the paper. Investigation: LR, RH. Funding acquisition, resources and supervision: MP, CAJ, SE.

## Acknowledgments

We would like to acknowledge funding from a Medical Research Council (MRC) research grant MR/M000532/1 (to CAJ), a Sir Jules Thorn Biomedical Award JTA/09 (to CAJ). RH was funded by an MRC PhD studentship award (1233630). LR was funded by a Biotechnology and Biological Sciences Research Council (BBSRC) White Rose Doctoral Training Partnership PhD studentship (BB/M011151/1).

