## Supplementary table and figures for "The role of MEGF10 in myoblast fusion and hypertrophic response to overload of skeletal muscle"

**In vivo and in vitro analyses of MEGF10 function demonstrate its role in myoblast fusion and hypertrophic response to overload of skeletal muscle**

**Louise Richardson, Ruth Hughes, Colin A Johnson, Stuart Egginton, Michelle Peckham**

**Supplementary Material**

**Supplementary Tables**

**Supplemental Table 1: Primary Antibodies used:**

| Antibody | Source | Species | Dilution (IF) | Dilution (WB) |
|---|---|---|---|---|
| A4.1025 | ATCC hybridoma | Mouse | 1:10 | N/A |
| c-myc (9E10) | Sigma | Mouse | N/A | 1:5000 |
| GFP (AB10145) | Millipore | Rabbit | N/A | 1:1000 |
| Pax7 supernatant. | DSHB (hybridoma) | Mouse | 1:20 | N/A |
| MyoD (MA1-41017) | Thermo Scientific | Mouse | 1:100 | 1:500 |
| MyoD (C-20) | Santa Cruz (sc-304) | Rabbit | 1:2000 | N/A |
| Myogenin (F5D) | DSHB | Mouse | 1:50 | 1:1000 |
| Myogenin (MA511486) | Invitrogen | Mouse | 1:50 | N/A |
| Laminin L9393 | Sigma | Rabbit | 1:30 | N/A |
| DAPI 40043 | Sigma | N/A | 1:500 | N/A |

**Supplemental Table 2. Genotypic ratio of mice born from Megf10+/- matings.** Total number of WT, heterozygous and homozygous mice and corresponding ratio.

| | **WT** | **Heterozygous** | **Homozygous** |
|---|---|---|---|
| **Total** | 13 | 20 | 6 |
| **Ratio** | 2 | 3 | 1 |

**Supplementary figures**

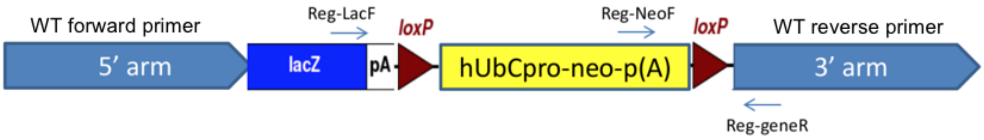


**Supplementary Fig. 1. Primer strategy for TaqMan assay for genotyping mice.** Schematic showing forward and reverse primer, human ubiquitin C gene promoter (hUbCpro-neo-p(A)) selection cassette flanked by loxP sites and lacZ sequence within the mutant (tm1) allele. Primers used in genotyping used sequence from lacZ (Reg-LacF) and the neomycin selection cassette (tm1a allele: Reg Neo-F), the reverse primer Reg-geneR. The readout of LAC Z+ WT+ = heterozygous, LAC Z+ WT- = homozygous, and LAC Z- WT+ = wild-type.

Sequences of the genotype primers were: LAC-Z: Forward primer: CGATCGTAATCACCCGAGTGT; Reverse primer: CCGTGGCCTGACTCATTCC, Reporter 1: CCAGCGACCAGATGAT; Reporter 2: N/A

And for MEGF10-1 WT; Forward primer: CTACCGGACAGCCTACCG; Reverse primer: CTTTCATAAAATCCTGGGCAACACT; Reporter 1: TATAGACGCAAATCCC, Reporter 2: N/A

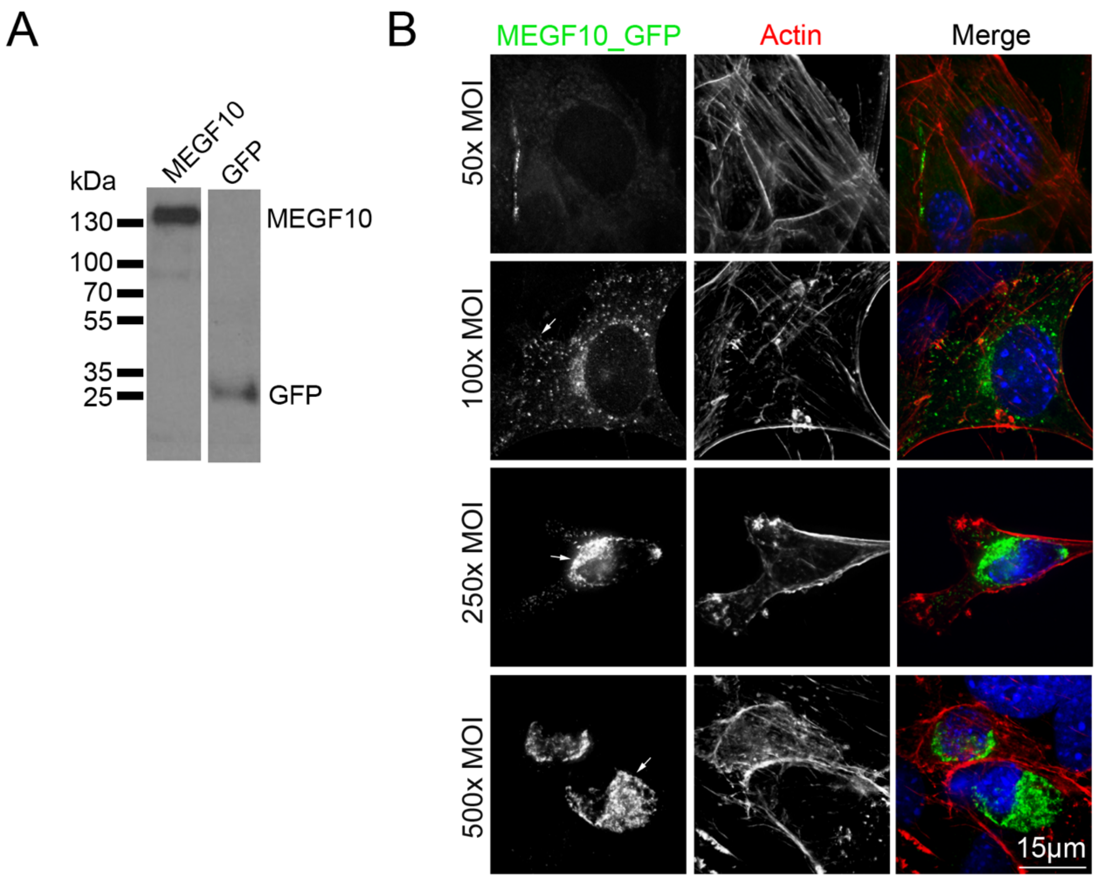


**Supplemental Fig. 2**: **eGFP and eGFP-MEGF10 expression tests.** **A**: anti-GFP blots for MEGF10-eGFP and the GFP adenovirus expressed in cells. **B:** Localisation of MEGF10-EGFP at different MOIs. At an MOI of 50, very little signal is observed. At an MOI of 100, MEGF10-eGFP is found in vesicles and at the plasma membrane (arrowed). At MOIs of 250 and 500, most of the MEGF10-EGFP is in the Golgi (arrowed)

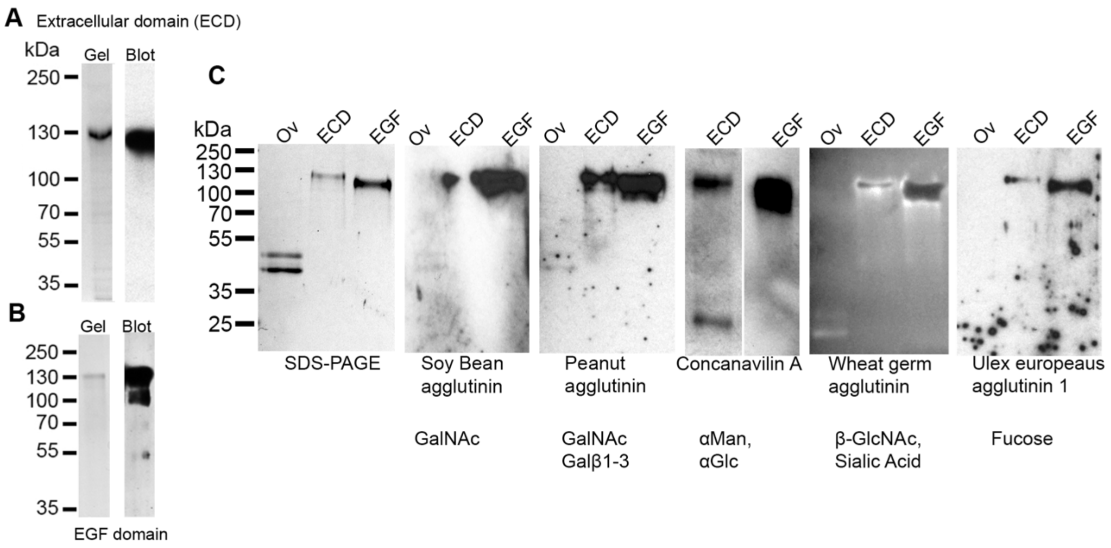


**Supplemental Fig 3: Expression, purification and Lectin blots for ECD and EGF. A and B:** Gels and blots (anti-Myc Tag) for purified extracellular domain (ECD) (**A**) and EGF domains (**B**) of MEGF10 to show the purified protein and any contaminants. **C**: SDS Page gels of Ovalbumin (Ov), ECD and EGF domains together with lectin blots using each of the specific lectins, and what they recognise, as shown.
